# Detailed measurements and simulations of electric field distribution of two TMS coils cleared for obsessive compulsive disorder in the brain and in specific regions associated with OCD

**DOI:** 10.1101/2022.01.13.476243

**Authors:** Marietta Tzirini, Yiftach Roth, Tal Harmelech, Samuel Zibman, Gaby S Pell, Vasilios k. Kimiskidis, Aron Tendler, Abraham Zangen, Theodoros Samaras

## Abstract

The FDA cleared deep transcranial magnetic stimulation (Deep TMS) with the H7 coil for obsessive-compulsive disorder (OCD) treatment, following a double-blinded placebo-controlled multicenter trial. Two years later the FDA cleared TMS with the D-B80 coil on the basis of substantial equivalence. In order to investigate the induced electric field characteristics of the two coils, these were placed at the treatment position for OCD over the prefrontal cortex of a head phantom, and the field distribution was measured. Additionally, numerical simulations were performed in eight Population Head Model repository models with two sets of conductivity values and three Virtual Population anatomical head models and their homogeneous versions. The H7 was found to induce significantly higher maximal electric fields (p<0.0001, t=11.08) and to stimulate two to five times larger volumes in the brain (p<0.0001, t=6.71). The rate of decay of electric field with distance is significantly slower for the H7 coil (p < 0.0001, Wilcoxon matched-pairs test). The field at the scalp is 306% of the field at a 3 cm depth with the D-B80, and 155% with the H7 coil. The H7 induces significantly higher intensities in broader volumes within the brain and in specific brain regions known to be implicated in OCD (dorsal anterior cingulate cortex (dACC), dorsolateral prefrontal cortex (dlPFC), inferior frontal gyrus (IFG), orbitofrontal cortex (OFC) and pre-supplementary motor area (pre-SMA)) compared to the D-B80. Significant field ≥ 80 V/m is induced by the H7 (D-B80) in 15% (1%) of the dACC, 78% (29%) of the pre-SMA, 50% (20%) of the dlPFC, 30% (12%) of the OFC and 15% (1%) of the IFG. Considering the substantial differences between the two coils, the clinical efficacy in OCD should be tested and verified separately for each coil.

## Introduction

Obsessive-compulsive disorder (OCD) is a chronic, disabling mental disorder with a lifetime prevalence of 2-3% [1]. Clinically, it is characterized by recurrent intrusive thoughts or obsessions and compulsive behaviors. Obsessions and compulsions cause distress and are mitigated by avoidance of situations believed to exacerbate the symptoms. The therapeutic mainstay involves high doses of serotonin reuptake inhibitors and/or exposure and response prevention therapy. Unfortunately, about half the patients do not benefit or remain significantly symptomatic [2], often due to intolerable side-effects [3–4]. Until recently, treatment options for such patients included switching to another serotonin reuptake inhibitor (SRI), augmenting with an antipsychotic, and/or treatment with more intensive cognitive behavioral therapy (CBT). Yet, since OCD is a chronic illness that frequently begins in adolescence [5], many patients will undergo various treatment trials during their lifetime without sufficient improvement [6–7]. Therefore, novel treatment approaches are of great need.

The neuroanatomical basis of dysfunction in OCD has been well known for many years and is localized to the cortico-striato-thalamic-cortical (CSTC) circuitry. Neuroimaging studies have disclosed aberrant patterns of connectivity amongst key nodes of the CSTC circuitry (e.g. [8–9]). Rarely, in extreme cases, patients with treatment refractory OCD undergo psychosurgery to certain elements of this circuit. In August 2018, the US FDA cleared deep repetitive transcranial magnetic stimulation (Deep TMS) with the H7 coil (BrainsWay Ltd, Israel) for the therapeutic management of OCD. The clearance was based on the results of a pilot sham-controlled study [10] and a subsequent sham-controlled multicenter study [11], where each Deep TMS session was preceded by an individually tailored provocation in order to activate the relevant circuitry and make it prone to change by the stimulation [12]. In the multi-center study [11] 38.1% of OCD patients reached response after six weeks of Deep TMS, and 45.2% were responders at the one-month follow-up. In August 2020, MagVenture (Denmark) obtained a US FDA 510K clearance, or statement of substantial equivalence, for their cooled 120° bent figure-8 coil, the D-B80 coil, based on electrical field modeling. The two coils differ in diameter of their circular wings and distance between the two wings. In addition, the D-B80 has a rigid fixed 120° angle between the two wings, while the H7 has flexible windings that conform to the subject’s head. We performed analysis of the electric field (EF) distributions of the two coils using both measurements and simulations. A comparison of the volume and depth of stimulation in the whole brain found significant differences between the two coils [13]. The present work includes extensive analysis of the EF distribution in the brain based on head phantom field measurements as well as simulations in 22 head models including colored field maps, histograms showing the distribution of field intensities in multiple models and EF penetration depth within the brain. Moreover, we investigate in detail the effects of the D-B80 and H7 coils in specific parts of the CSTC circuitry known to be associated with the pathophysiology of OCD and discuss connections between these effects and clinical results found in previous studies.

## Materials and Methods

### Electric field measurements in head phantom

For the H7 and D-B80 coils, the induced EF characteristics, including amplitude and orientation, were measured in an experimental phantom of the human head (13 cm × 18 cm, height 15 cm) filled with physiologic saline solution. The field was measured using a two-wire dipole probe [14] with 1.5 cm distance between the two tips. The probe was moved in three directions inside the head phantom using a displacement system with 0.5 mm resolution, and the field distribution of each coil was measured in the whole head phantom volume with 1 cm resolution. The coil location was over the prefrontal cortex (PFC), at the treatment position according to the FDA-cleared protocol, which is 4 cm anterior to the leg motor cortex [10–11]. At each measurement point the field was measured along two perpendicular directions, i.e., along the posterior-anterior axis and along the lateral-medial axis. At the PFC location, the coils induce EF also along the superior-inferior axis. Hence, the coils were also attached, at the same prefrontal location, to a head phantom lying with the forehead down, so that the probe could measure the lateral-medial and superior-inferior axes. The maps from the two phantoms were combined to create 3D maps for each of the coils. The probe was connected to a high impedance PC-based oscilloscope (Agilent, USA), and the measured maximal voltage divided by the distance between the wire tips gave the EF along each axis. Pulses were recorded using in-house software. The absolute magnitude and the direction of the EF were computed for each measurement point.

Maps of EF distribution based on these measurements were produced using MATLAB (MATLAB R2019a, The MathWorks Inc., Natick, Massachusetts) for each coil.

The procedure for treatment of OCD according to the cleared protocol includes determination of the leg resting motor threshold (rMT), and then the treatment is given at 100% of the leg rMT. The threshold was set to 100 V/m, which is within the accepted range of thresholds required for motor activation [15–19], at a depth of 3 cm, which is considered the average depth of the leg motor cortex representation [20–21]. The D-B80 was connected to a MagPro R30 stimulator (MagVenture A/S, Denmark), with a maximal output of 1.8 kV. The H7 was connected to a BrainsWay stimulator (BrainsWay Ltd, Israel), with a maximal output of 1.7 kV. The maps were adjusted to the average percentage of the maximal stimulator output (MSO) empirically found to achieve 100% of the leg rMT. This was 53% of MSO for the D-B80, based on previous studies [22–23], and 54% for the H7 ([10–11], BrainsWay data on file). At this power output, the coil currents were 3.64 kA for the D-B80 and 3.18 kA for the H7.

The distribution of values of EF intensity induced in the brain by each coil, were plotted as histograms, with volumes in cm^3^ for each field range.

### Electric field simulations in head models

Both coils were simulated as current sources, with a frequency of 3.5 kHz and a current amplitude of 3.64 kA for the D-B80 and 3.18 kA for the H7, in the Sim4Life (S4L) platform (Sim4Life 6.0, Zurich MedTech AG, Switzerland). The D-B80 model, generated by Makarov and colleagues [24], consists of 2 layers including 3 windings on top and 4 windings beneath with outer and inner diameter of 95 mm and 67 mm, respectively. The H7 model, based on a CAD file provided by the manufacturer, had 2 layers of 4 elliptically shaped windings each, one on top of the other, whose major axis ranged from 130 to 70 mm and minor axis from 105 to 55 mm. The coils were placed over the PFC of three high resolution anatomical models of the Virtual Population (ViP) family (Duke, Ella, Thelonious) [25–26], and eight members of the Population Head Model (PHM) repository [27]. For better comparison with the electric field measurements inside the phantom, homogeneous versions of the three ViP models were also used. All models, except Thelonious, which is a model of a 6-year-old child, are adults aged from 22 to 35 years old. To evaluate the impact of conductivity variations in literature, two sets of conductivity values were used for the tissues of the PHM repository models. The first set (*PHM v.1*) corresponds to the low frequency tissue database of IT’IS Foundation [28], while the second (*PHM v.2*) includes typical conductivity values for TMS simulations [29] (S1_Table).

Both coils were placed as close as possible to each head. Moreover, since the H7 is flexible, the angle of the coil was fitted individually for each model, in such a way that its elements were fully tangential to the head.

The heads of ViP models in their original version include more than 30 different structures, which were all assigned to the same electrical conductivity value (1.2 S/m) for their homogeneous version. The volume denoted as brain in ViP models consists of 14 structures, including white matter, grey matter, cerebellum, cerebrospinal fluid (CSF), corpus callosum, anterior and posterior commissure, hippocampus, hypophysis, hypothalamus, medulla oblongata, pineal body, pons, and thalamus. The heads of PHM repository models consist of ventricles, white matter, grey matter, cerebellum, CSF, skull, and skin. White matter, grey matter and cerebellum were denoted as brain.

The EF distributions were computed by S4L’s magneto-quasi-static (M-QS) solver and then compared, using MATLAB scripts, in terms of maximum value (E_max_), maximal depth (d_100_) for which EF ≥ 100 V/m and stimulated volume (V_100_) in the brain. The 99.5^th^ percentile of EF distribution was considered as E_max_ to avoid numerical artifacts. The part of volume where the EF was greater than 100 V/m was defined as V_100_.

The distribution of values of EF within the brain and specific brain regions which comprise parts of the CSTC circuitry were quantified as histograms in bins of 10 V/m. The specific regions were the pre-supplementary motor area (pre-SMA), inferior frontal gyrus (IFG), dorsolateral prefrontal cortex (dlPFC), orbitofrontal cortex (OFC), and dorsal anterior cingulate cortex (dACC). In these brain regions, the descriptive statistics of the EF amplitude distribution (i.e., 25th, 50th, 75th, and 99th percentiles of the distribution) of both coils were quantified for each of the 22 head models. In addition, the percentage of each region volume with significant induced EF ≥ 80 V/m were calculated in each head model, and the descriptive statistics across models was quantified. The Brodmann atlas template of MRIcron tool (MRIcron v1.0.20190902, McCausland Center for Brain Imaging, University of South Carolina) was used to locate the above brain regions. The atlas image and the voxelized data of each PHM model were co-registered through Statistical Parametric Mapping (SPM) software (SPM12, The Wellcome Centre for Human Neuroimaging, UCL Queen Square Institute of Neurology, London), to identify pre-SMA as Brodmann area (BA) 8, IFG as BA 44 and 45, dlPFC as BA 9 and 46, OFC as BA 10, 11 and 47 and dACC as BA 24, 25 and 32. Additionally, a 3D segment of the atlas was manually applied to the brain of each ViP model to extract the corresponding masks.

### Statistical Methods

The stimulation volumes and depths and the maximal EF values within the brain were compared using paired t-test. The rates of decay of EF with distance and the distribution of values of EF within the brain and specific regions were compared using Wilcoxon matched-pairs test (after failed normality test). The results were considered significant if p < 0.05.

## Results

### Results of electric field measurements in head phantom

Fig 1 illustrates color intensity maps of the EF distribution induced in the brain by the two coils located at the treatment location over the PFC, superimposed on brain coronal slices of T1-weighted anatomical MR images. The areas in red indicate field magnitude at or above 100 V/m.

**Fig 1.**
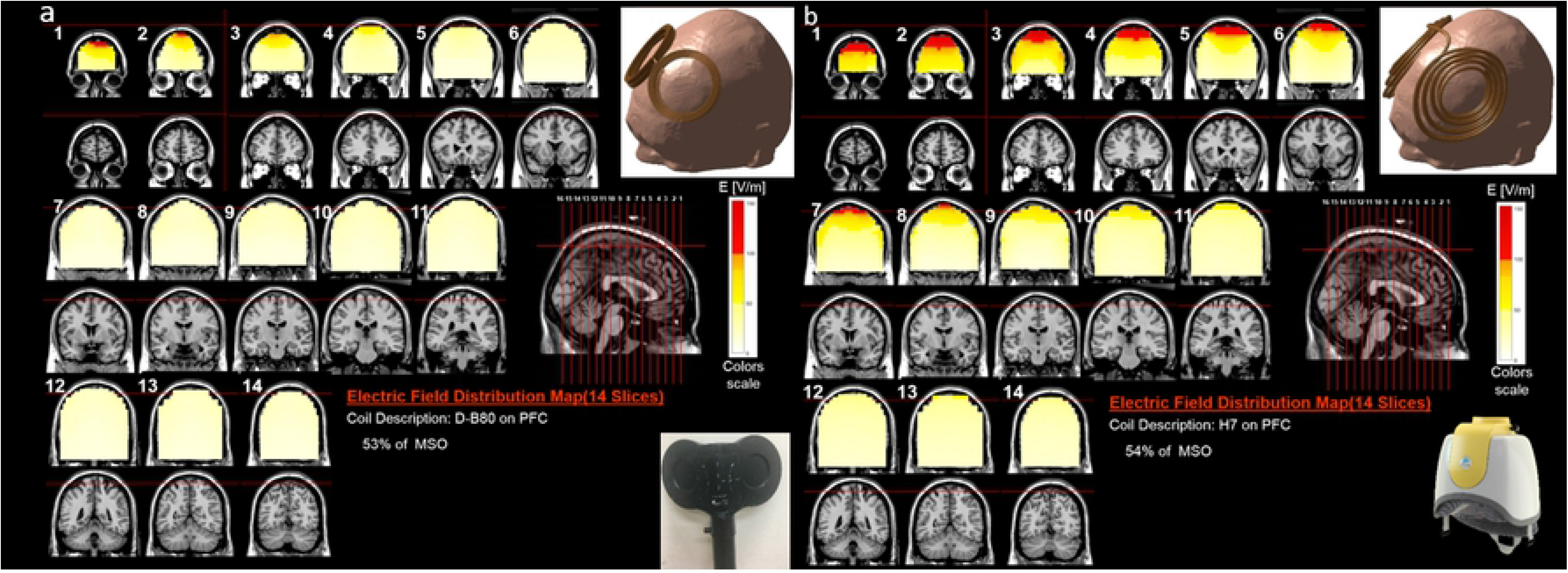
Colored maps of electric field distribution. Colored field maps for the D-B80 (a) and for the H7 (b) showing electric field distribution within the brain, when located at the treatment location over the prefrontal cortex, indicating the electric field absolute magnitude in each pixel over 14 coronal slices 1 cm apart. The maps were adjusted to the average percentage of the maximal stimulator output required to achieve 100% of the leg rMT, which are 53% for the D-B80 and 54% for the H7. The red pixels indicate field magnitude ≥ the threshold for neuronal activation, which was set to 100 V/m.

As can be seen in Fig 1, when placed over the PFC, the H7 stimulates much larger and deeper brain volume (more red pixels), and induces higher field intensities, while the effect of the D-B80 at this location is much weaker, and supra-threshold field is induced only in very shallow brain layers. The distribution of values of EF intensity within the brain are plotted as histograms for the two coils in Fig 2, showing the volume with induced field in each field range.

**Fig 2.**
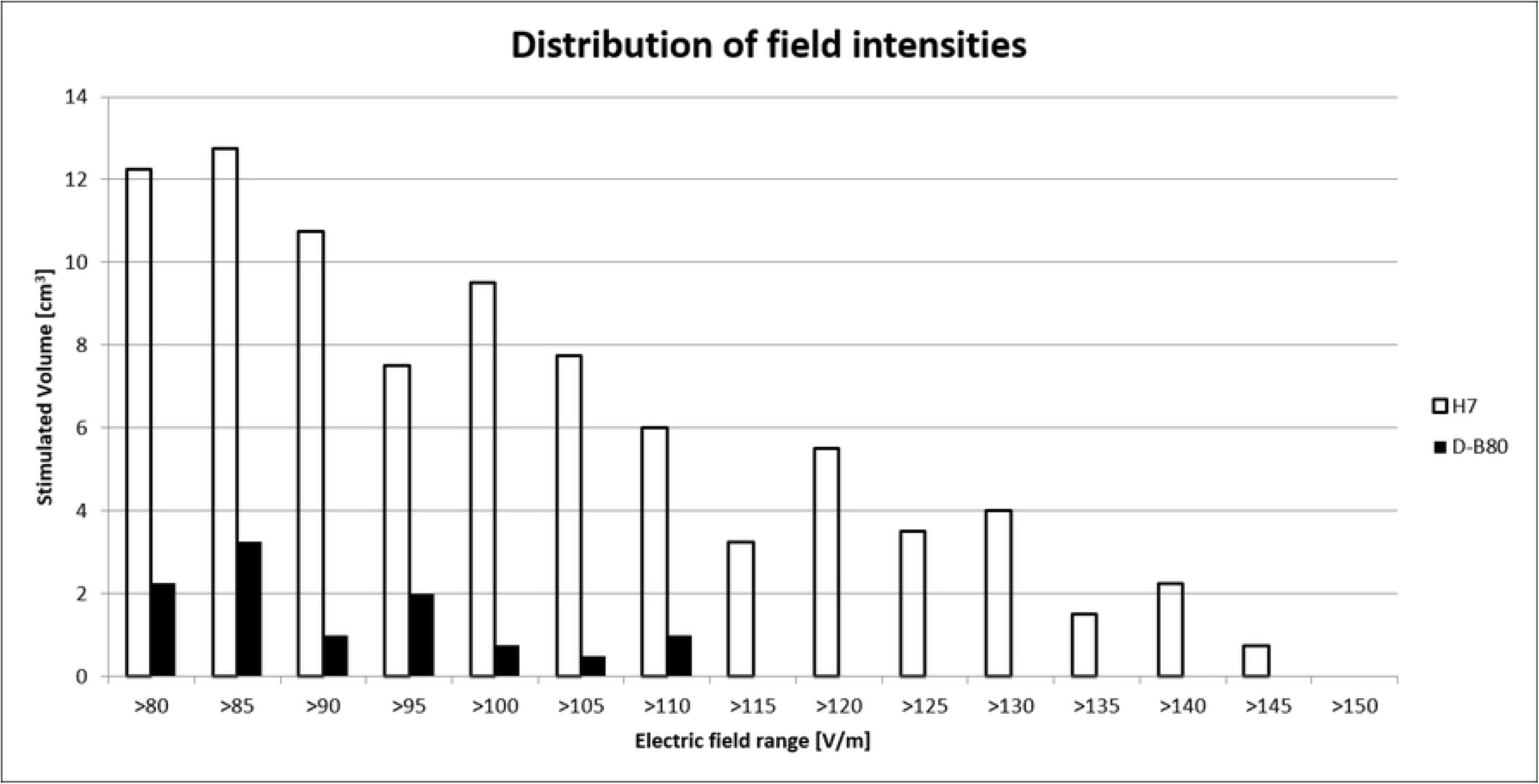
Distribution of values of EF intensity. Histograms of distribution of the volume in cm^3^ according to the induced electric field range within the brain for the D-B80 and H7. Field columns are in bins of 5 V/m.

The distribution of field values within the brain is significantly different between the coils (p=0.002, Wilcoxon matched-pairs test).

The EF decay profile as a function of distance from the coil when located over the PFC is shown in Fig 3 for both coils, normalized to the field at the scalp, 0.5 cm from coil surface (Fig 3b), to the field at the brain surface, 1.5 cm from coil surface (Fig 3c), and to the field at a depth of 3 cm from the coil (Fig 3d). The field was measured along a line in a central sagittal plane starting at the point of inflection of the frontal bone and going at 45° downward and posteriorly (Fig 3a).

**Fig 3.**
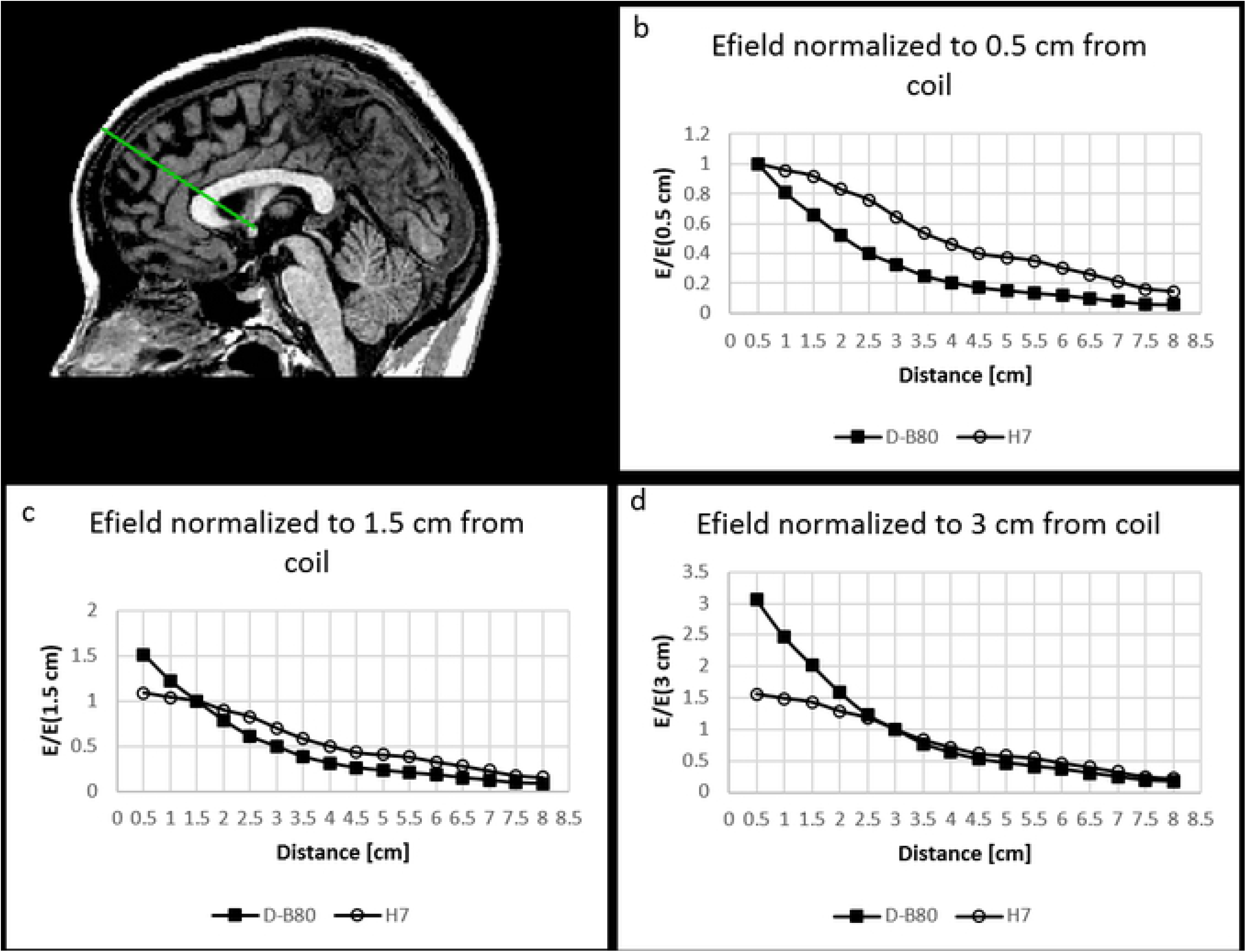
Field decay profile. The electric field magnitude measured in a phantom head model as a function of the distance from the coil, is shown for the D-B80 coil and for the H7 coil, when located at the treatment location over the prefrontal cortex. The field was measured along a line in a central sagittal plane starting at the point of inflection of the frontal bone and going at 45° downward and posteriorly (a). The electric field was normalized to the field at the scalp, 0.5 cm from coil surface (b), to the field at the brain surface, 1.5 cm from coil surface (c), and to the field at a depth of 3 cm from coil surface (d).

The rate of decay of EF with distance is significantly slower for the H7 coil compared to the D-B80 coil, when located at the treatment location over the PFC (p < 0.0001, Wilcoxon matched-pairs test). The field at the scalp is 306% of the field at a 3 cm depth with the D-B80, and 155% with the H7 coil (Fig 3d). The field at the scalp is 152% of the field at the brain surface with the D-B80, and 109% with the H7 coil (Fig 3c).

### Results of electric field simulations in head models

The results of E_max_, d_100_ and V_100_ for all 22 models, as well as the results of the phantom head model measurements, are shown in Table 1.

**Table 1.** Stimulation volumes V_100_ (cm^3^), stimulated depth d_100_ (mm) and maximal electric field values E_max_ (V/m) in the brain for simulations in 22 models and for phantom measurements. Shown are results of comparison of stimulation volumes V_100_ in the brain between the D-B80 and H7 coils using Wilcoxon matched-pairs test.

| Model | | $E_{\max}$ (99.5th) [V/m] | | $d_{100}$ [mm] | | $V_{100}$ [cm<sup>3</sup>] | | Wilcoxon matched-pairs test |
| --- | --- | --- | --- | --- | --- | --- | --- | --- |
|  |  | D-B80 | H7 | D-B80 | H7 | D-B80 | H7 | p-value |
| Homogeneous | Thelonious | 135.7 | 159.4 | 15.31 | 20.23 | 23.6 | 93.4 | 0.0005 |
|  | Ella | 106.4 | 127 | 10.92 | 14.98 | 8.6 | 31 | 0.0313 |
|  | Duke | 100.5 | 113.6 | 9.81 | 11.08 | 6.6 | 22.6 | 0.25 |
| ViP | Thelonious | 148.4 | 189.8 | 41.69 | 51.82 | 28.9 | 122.3 | 0.0002 |
|  | Ella | 113.8 | 142.2 | 55.64 | 61.11 | 11.9 | 38.3 | 0.0039 |
|  | Duke | 118.3 | 140.4 | 36.68 | 44.49 | 14.3 | 46 | 0.0039 |
| PHM v.1 | #101309 | 118.6 | 123.4 | 16.59 | 26.76 | 13.4 | 20.9 | 0.0625 |
|  | #103111 | 109.7 | 123.7 | 18.91 | 20.04 | 9.7 | 20.2 | 0.0625 |
|  | #103414 | 107.2 | 144.4 | 15.02 | 25.54 | 7.8 | 38.7 | 0.0039 |
|  | #105014 | 100.6 | 127.9 | 16.29 | 24.2 | 6 | 23.4 | 0.0313 |
|  | #105115 | 108.8 | 139.6 | 14.25 | 32.05 | 8.9 | 35.6 | 0.0078 |
|  | #106016 | 96.9 | 133.5 | 15.99 | 22.96 | 4.7 | 24.5 | 0.0156 |
|  | #110411 | 120.9 | 147.6 | 14.73 | 33.67 | 14.6 | 51.1 | 0.002 |
|  | #111716 | 113.1 | 127.1 | 11.22 | 30.5 | 9.6 | 26.3 | 0.0313 |
| PHM v.2 | #101309 | 122.5 | 129.4 | 12.52 | 24.08 | 14.5 | 24.7 | 0.0313 |
|  | #103111 | 112.1 | 128.3 | 14.93 | 40.04 | 10.6 | 23.4 | 0.0313 |
|  | #103414 | 109 | 149.2 | 12.86 | 34.89 | 8.4 | 44.4 | 0.002 |
|  | #105014 | 102.9 | 133.9 | 16.9 | 26.43 | 6.6 | 27 | 0.0156 |
|  | #105115 | 109.8 | 143.8 | 16.17 | 44.94 | 9.3 | 39.5 | 0.0039 |
|  | #106016 | 99.2 | 138.3 | 19.55 | 23.51 | 5.3 | 28 | 0.0078 |
|  | #110411 | 123.7 | 155.1 | 17.04 | 58.3 | 15.8 | 59.2 | 0.0005 |
|  | #111716 | 117.3 | 137.6 | 21.55 | 30.53 | 11.5 | 34.9 | 0.0078 |

| Model |  | E<sub>max</sub> (99.5th) [V/m] |  | d<sub>100</sub> [mm] |  | V<sub>100</sub> [cm<sup>3</sup>] |  | Wilcoxon<br>matched-pairs test |
| --- | --- | --- | --- | --- | --- | --- | --- | --- |
|  |  | D-B80 | H7 | D-B80 | H7 | D-B80 | H7 | p-value |
| Experimental | Phantom | 120 | 152 | 10 | 20 | 4.3 | 52 | 0.002 |
| Mean |  | 113.7 | 139.4 | 18.9 | 31.39783 | 11.1 | 40.3 |  |
| SD |  | 12 | 15.8 | 11.0 | 13.24197 | 5.9 | 24.3 |  |

Comparison of the results shows that the H7 induces significantly higher maximal values in the brain of all models (p<0.0001, t=11.08), higher d_100_ (p < 0.0001, t =6.17) and stimulates two to five times larger volumes in the brain (p<0.0001, t=6.71).

Color intensity maps of the EF distribution induced in the brain by the two coils located at the treatment location over the PFC, for the 14 simulated head models, in the 6 most frontal coronal slices 1 cm apart, are shown in S1 Fig. The field maps for the 8 PHM v.2 models are plotted in S2 Fig.

The distributions of values of EF intensity within the brain are plotted as histograms in Fig 4 for the 14 numerical head models. The distributions of values for the 8 PHM v.2 models are plotted in S3_Fig.

**Fig 4.**
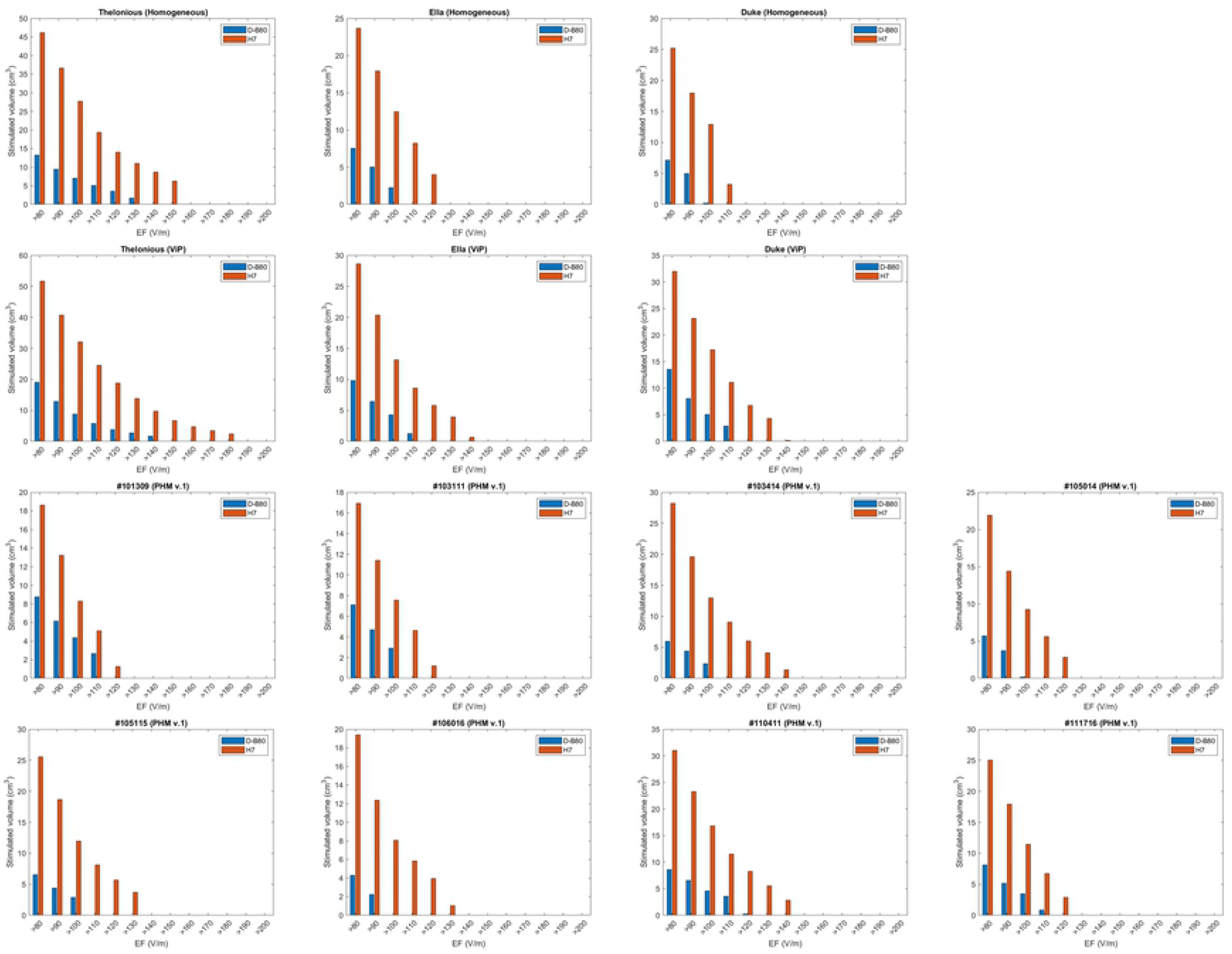
Distribution of values of EF intensity. Histograms of distribution of volume in cm^3^ within the brain according to the induced electric field range for the D-B80 and H7 coils. Field columns are in bins of 10 V/m. Results are shown for the 14 numerical head models.

The EF decay profile as a function of distance from the coil, normalized to the field at a depth of 3 cm from the coil, is shown in Fig 5 for the 14 numerical head models. The figures show an excellent agreement between the calculated decay profile inside homogeneous numerical models and experimental phantoms. The field decay profiles for the 8 PHM v.2 models are plotted in S4_Fig.

**Fig 5.**
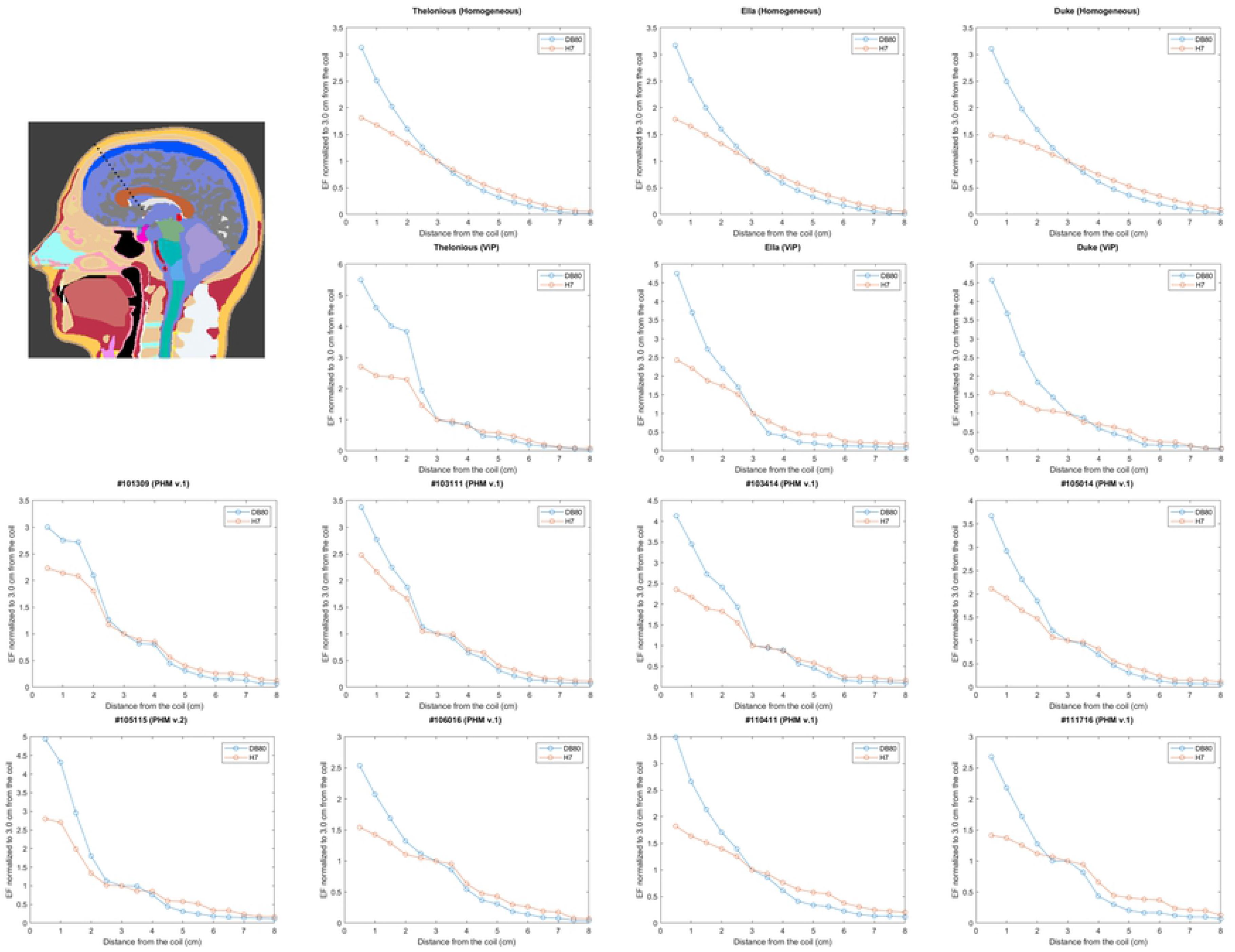
Field decay profile. The electric field magnitude as a function of the distance from the coil, is shown for the D-B80 coil and for the H7 coil, when located at the treatment location over the prefrontal cortex. The field was measured along a line in a central sagittal plane starting at the point of inflection of the frontal bone and going at 45° downward and posteriorly (top left). The electric field was normalized to the field at a depth of 3 cm from coil surface. Shown results for the 14 numerical head models.

Comparison results (Wilcoxon matched-pairs test) of the EF decay profile with distance are shown for the 22 numerical head models in S2_Table.

The distribution of values of EF intensity within five brain regions (dACC, dlPFC, IFG, OFC and pre-SMA) are plotted as histograms in Fig 6, depicting the averaged results of all the 22 head models. Comparison of the EF distribution using Wilcoxon matched-paired test found a very significant difference between the coils of p<0.0001 in all five brain regions.

**Fig 6.**
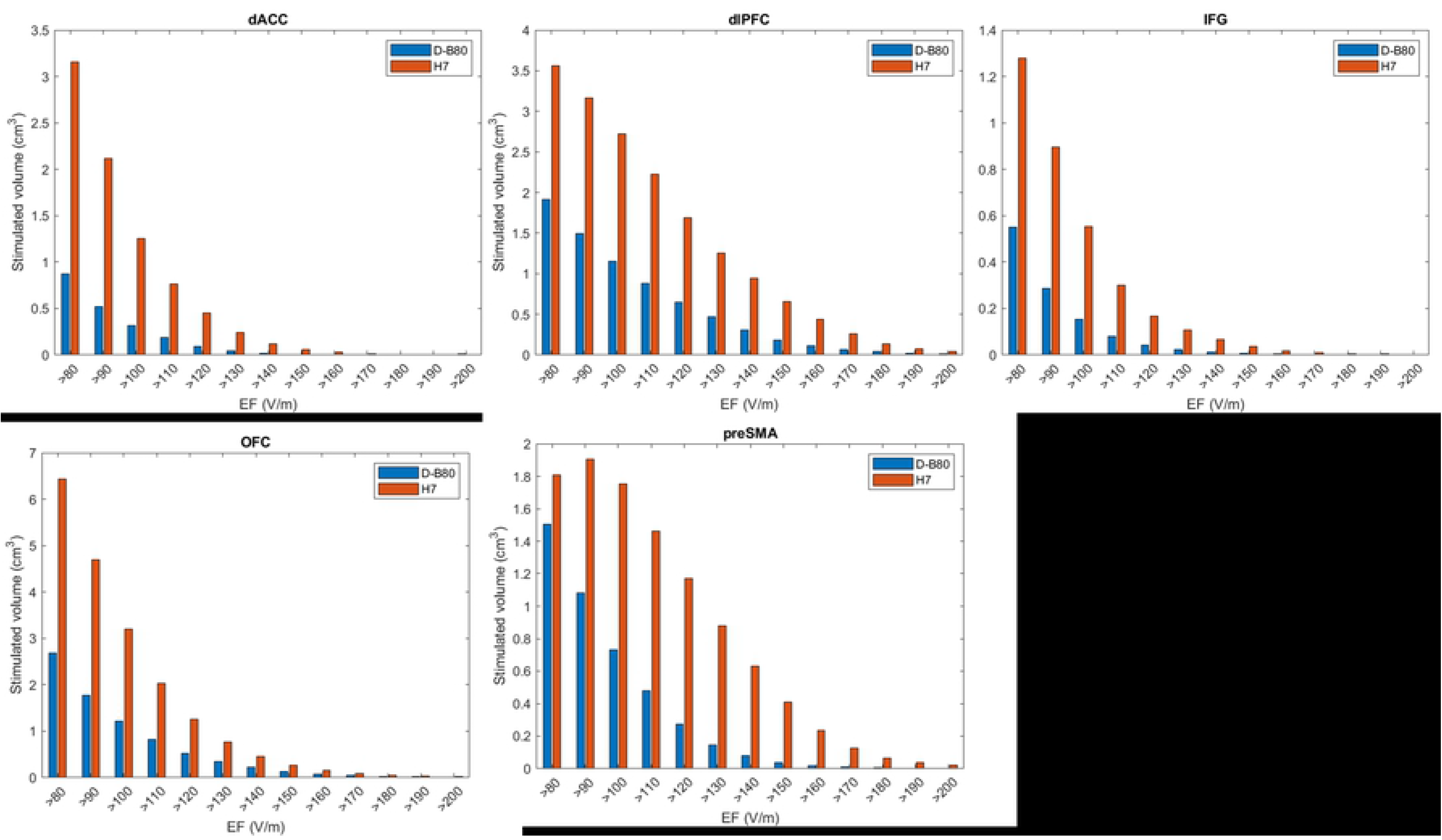
Distribution of values of EF intensity within specific brain regions. Histograms of distribution of the volume in cm^3^ according to the induced electric field range for the D-B80 and H7 coils, are plotted for five brain regions (dACC, dlPFC, IFG, OFC and pre-SMA). Field columns are in bins of 10 V/m. Shown the averaged results of all the 22 simulated head models.

Fig 7 illustrates for each of the five brain regions the descriptive statistics (25th, 50th, 75th, and 99th percentile) of the percentage of region volume with significant induced EF ≥80 V/m for the D-B80 (a) and H7 (b) coils, based on the 22 head models. In addition, the descriptive statistics of the EF amplitude distributions within these regions are shown for the D-B80 (c) and H7 (d) coils.

**Fig 7.**
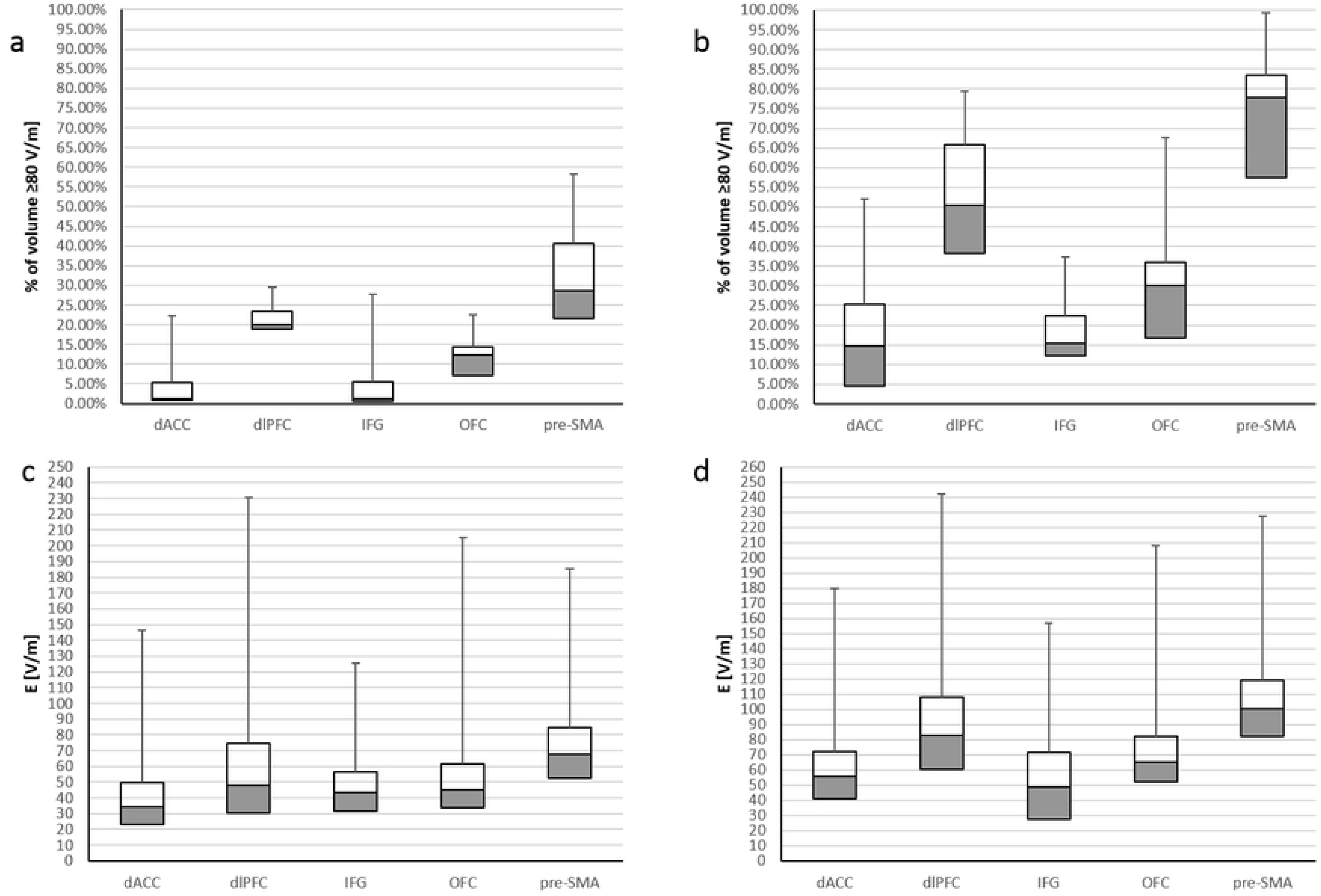
Descriptive statistics within specific brain regions. Descriptive statistics across 22 head models for five brain regions (dACC, dlPFC, IFG, OFC and pre-SMA) of the percentage of EF ≥80 V/m for the D-B80 (a) and the H7 (b), and of the EF amplitude distribution within these regions for the D-B80 (c) and the H7 (d). dACC: dorsal anterior cingulate cortex; dlPFC: dorsolateral prefrontal cortex; IFG: inferior frontal gyrus; OFC: orbitofrontal cortex; pre-SMA: pre-supplementary motor area.

From this figure, one can see that the H7 induces EF ≥80 V/m in 15% (5% to 25%) (median (25th to 75th percentiles) across the 22 models) of the dACC, 50% (38% to 66%) of the dlPFC, 15% (12% to 22%) of the IFG, 30% (17% to 36%) of the OFC and 78% (57% to 83%) of the pre-SMA. The D-B80 induces EF ≥80 V/m in 1.3% (0.8% to 5.2%) of the dACC, 20% (19% to 24%) of the dlPFC, 1.3% (0.6% to 5.6%) of the IFG, 12% (7% to 14%) of the OFC and 29% (22% to 41%) of the pre-SMA. The median EF induced by the H7 is 100 V/m in the pre-SMA, 83 V/m in the dlPFC, 65 V/m in the OFC, 56 V/m in the dACC and 49 V/m in the IFG. The D-B80 induces median EF of 68 V/m in the pre-SMA, 34 V/m in the dACC and below 50 V/m in the other three regions.

## Discussion

In this study we did a comprehensive analysis of the EF distribution of the two TMS coils that the FDA has cleared for OCD treatment, with simulations in 22 different numerical head models, as well as EF measurements in a head phantom with the actual coils. In all models the coils were placed at the treatment location, over the medial PFC (mPFC). There is high variability in the results between the various models, most probably due to the large differences in size and internal compartmentation. Yet, all the methods and models found that the H7 stimulates significantly broader and deeper brain volume compared to the D-B80. The rate of field attenuation as a function of distance from the coil is significantly slower for the H7. This means that at the power output used for treatment (100% of the foot motor threshold), many deeper structures are stimulated by the H7 but not by the D-B80. Alternatively, should a higher power output be used with the D-B80 in order to stimulate deeper structures, the induced field at the brain surface will be higher. The aggregate of results clearly indicates that many prefrontal structures are stimulated by the H7 but not by the D-B80. Among those are structures within the pre-SMA, IFG, dlPFC, OFC and the dACC (Figs 6 and 7). All of these comprise parts of the CSTC circuitry and as such, have been implicated to different extents in the pathophysiology of OCD [30–32].

OCD is a complex and heterogeneous disorder associated with high personal and societal costs. Deficits in cognitive and affective processing in OCD patients are mediated by alterations in specific neural circuits [32]. Data from functional and structural imaging studies support this hypothesis by demonstrating alterations in several brain regions that that are involved in sensorimotor, cognitive, affective, and motivational processes of OCD patients compared to healthy individuals [32]. The pre-SMA and IFG are key components of the CSTC that are involved in response inhibition [33–37]. While decreased gray matter volume and decreased task-related activation during inhibition was found in the IFG of OCD patients [38–40], the pre-SMA has demonstrated hyperactivation in both OCD patients and their unaffected siblings and is thought to be a compensatory mechanism [41]. Several studies noted that abnormalities in the dlPFC, a key structure involved in executive control, is closely related to symptomatic features of OCD, including excessive doubt and repetitive actions [42–45]. Task-based fMRI studies employing cognitive and executive tasks indicated decreased responsiveness of the dlPFC during task planning in OCD patients [46–47]. In the OFC, which is crucially involved in reward-guided learning and decision making [48], lower activity was identified in neuroimaging studies during reversal learning [49–50], and OFC hyperactivity was found at rest [51–52] and during OCD symptom provocation [53–56]. This hyperactivity normalizes with successful treatment [51, 57-59]. Other corticostriatal pathways, most prominently pathways through the dACC and dmPFC, were implicated in OCD pathology [60]. Feelings of doubt, worry, and repetitive behavior, key symptoms of OCD, have been linked to hyperactive error signals in the brain. The error-related negativity (ERN) [61] is an event-related potential in the theta frequency band (4-8 Hz) that peaks 50-150 milliseconds following errors on speeded reaction-time (RT) tasks (e.g., the Stroop, flanker). The ERN is consistently attributed to the dACC [62–63] and is thought to reflect processes that are involved in monitoring ongoing task performance, including those involved in making behavioral adjustments to prevent mistakes, the monitoring of cognitive conflict, or the emotional response to errors [61, 64-66]. In response to self-induced mistakes, there is over-activation of a specific monitoring system that is most pronounced in the dACC [64, 67-74]. For example, dACC hyperactivity has been consistently reported in OCD participants during tasks that include the commission of a mistake, such as Stop-Signal, Flanker, or Stroop [66-69, 75]. Increased ERN amplitudes and post-error slowing (PES) have also been robustly reported in OCD over the last 20 years (e.g. [76–78]). Notably, apart from being enhanced in OCD participants [61, 71, 74, 79-82], ERN was also found to be potentiated among their unaffected first-degree relatives [83–85]. In addition, activity following perceived mistakes (Perceived Error-Related Theta Activity (PERTA)) that shares the same scalp distribution (over the mPFC), brain source localization (dACC), and theta band frequency (4-8 Hz) as the ERN found for self-induced errors [85–87] was also found to be increased in OCD patients and their unaffected siblings [88]. This suggests that the constantly over-activated detection system in OCD evolves in vulnerable individuals with neuronal predisposition.

The most beneficial interventions in OCD have been those that were able to target the mPFC and dACC. Increased resting state and task-related activity in the dACC were found in participants that improved after CBT treatment. Modulation of dACC activity may be a primary mechanism of action of CBT for OCD [89]. TMS 1 Hz treatment over the mPFC with a double-cone coil was shown to improve both OCD symptoms and response time, again suggesting a correlation between error monitoring impairment and OCD pathophysiology [78]. Double-blinded randomized controlled trials [10–11] have demonstrated that high frequency (HF) Deep TMS with the H7 over the mPFC and dACC is a safe and effective intervention for the alleviation of OCD symptoms in participants who failed to receive sufficient benefit from previous treatments. It was found that, compared to sham treatment, the response rate following H7 treatment was significantly higher for up to one month, and that the reduction in symptoms severity was correlated with the magnitude of changes in the ERN response [10]. A recent ^1^H MRS study [90] found significant increases in levels of NAA, Choline and Creatine in the dACC following Deep TMS with the H7 coil in OCD patients, indicating direct neural stimulation in this region. Recently it has been demonstrated that in real-world clinical practice, Deep TMS with H7 over mPFC-dACC was beneficial for the majority (73%) of OCD patients with the onset of improvement usually occurring after 20 sessions [91]. Clinical experience with the DB-80 in OCD is minimal and currently consists of one open-label study in 20 OCD patients found that 50% responded to treatment [59]. To the best of our knowledge, no RCTs have been reported to date.

## Conclusions

This study used comprehensive simulations in 22 head models in addition to phantom field measurements with the actual coils and found consistently that the H7 stimulated broad and deep prefrontal brain regions and induced significant EF in large volumes within the dACC and other brain structures associated with OCD. These findings provide further insight to the neuroanatomical target of the H7 that has been shown to alleviate OCD symptoms in a multicenter sham-controlled trial. Consistently, substantial differences were found between the H7 and D-B80 coils in EF distribution and stimulated brain volume. The H7 stimulated significantly broader and deeper brain volumes compared to the D-B80. Due to the differences of the electric fields induced by TMS coils, which can be substantial, as clearly shown in this study for H7 and D-B80, the clinical efficacy of any coil approved for OCD treatment should be independently investigated in appropriately powered, randomized controlled trials.

## Supporting information

**S1 Table. Electrical conductivity (S/m) values used for the PHM models simulations.**

**S2 Table. Comparison of electric field decay profiles.** Results of comparison of the electric field decay profile with distance between the D-B80 and the H7 coils using Wilcoxon matched-paired test, for simulations in 22 head models.

**S1 Fig. Colored maps of electric field distribution.** Colored field maps for the D-B80 coil (top row) and for the H7 coil

(bottom row) showing electric field distribution within the brain, when located at the treatment location over the prefrontal cortex, indicating the electrical field absolute magnitude in each pixel over the 6 most frontal coronal slices 1 cm apart. The red pixels indicate field magnitude ≥ the threshold for neuronal activation, which was set to 100 V/m. Shown are results for the 14 numerical head models.

**S2 Fig. Colored maps of electric field distribution.** Colored field maps for the D-B80 coil (top row) and for the H7 coil (bottom row) showing electric field distribution within the brain, when located at the treatment location over the prefrontal cortex, indicating the electrical field absolute magnitude in each pixel over the 6 most frontal coronal slices 1 cm apart. The red pixels indicate field magnitude ≥ the threshold for neuronal activation, which was set to 100 V/m. Shown are results for the 8 PHM v.2 simulated head models.

**S3 Fig. Distribution of values of EF intensity.** Histograms of distribution of the number of pixels within the brain according to the induced electric field range for the D-B80 and the H7 coil. Field columns are in bins of 5 V/m. Shown results for the 8 PHM v.2 simulated head models.

**S4 Fig. Field decay profile.** The electric field magnitude as a function of the distance from the coil, is shown for the D-B80 coil and for the H7 coil, when located at the treatment location over the prefrontal cortex. The field was measured along a line in a central sagittal plane starting at the point of inflection of the frontal bone and going at 45° downward and posteriorly (top left). The electric field was normalized to the field at a depth of 3 cm from coil surface. Shown results for the 8 PHM v.2 simulated head models.

